# Expanding the Cell-Free Reporter Protein Toolbox by Employing a Split mNeonGreen System to Reduce Protein Synthesis Workload

**DOI:** 10.1101/2023.08.24.554711

**Authors:** Caroline E. Copeland, Chloe J. Heitmeier, Khoa D. Doan, Shea C. Lee, Yong-Chan Kwon

**Author notes:** Corresponding Author: E-mail address (Yong-Chan Kwon). These authors contributed equally (Caroline E. Copeland, Chloe J. Heitmeier).

## Abstract

The cell-free system offers potential advantages in biosensor applications, but their limited substrate supply poses a challenge in balancing enough sensing resources to detect low limits of analyte while providing a robust output signal. In this study, we harnessed split versions of fluorescent proteins, particularly split super-folder green fluorescent protein and mNeonGreen, to improve energy efficiency and enhance detection limit in the cell-free system. A comparative analysis of the expression of 1-10 and eleventh segments of beta strands in both whole-cell and cell-free platforms revealed distinct fluorescence patterns. Moreover, integrating SynZip peptide linkers substantially improved complementation, achieving a fluorescence intensity reaching 73.6% of the full-length protein and a 4.8-fold increase in expression compared to the split system without the SynZip peptide linkers. The split protein reporter system can enable energy-efficient sensing of low analyte levels in the cell-free system, broadening the toolbox of cell-free biosensor repertoire.

## Introduction

The cell-free system (CFS) has proved to be an ideal system for biosensor design and development, leveraging its open architecture, selectivity for a specific pathway of interest, and cellular pathway mimicry to facilitate precise sensing mechanisms.^1^ While cell-free biosensors harness complex gene circuits for refined biosensing, intrinsically limited substrate supply in the CFS causes consequent energy depletion.^2^ Therefore, larger reporter proteins that require more energy and nutrients will exhaust the system substrate faster, shifting energy distribution in the CFS system away from the sensing side of the circuit. In this context, we integrated the split fluorescent protein system into the CFS, aiming for a more energy-efficient biosensor reporter to conserve energy in the system and reach lower limits of detection by allowing more reporter protein to be made.

Split fluorescent proteins, such as super-folder green fluorescent protein (sfGFP)^3-5^ and mNeonGreen (mNG),^6,7^ fused with the proteins of interest, have previously been used to visualize protein-protein interactions. Typically, the split fluorescent protein application involves splitting the eleventh β-strand from the 11-segment β-stranded barrel of the green fluorescent protein. The reconstitution of fluorescent signal is spontaneous, without a covalent linkage, facilitated by its barrel-shaped protein’s tolerance to circular permutation.^8^ This study sought to express the small eleventh β-strand (17 amino acids long, 2.0 (sfGFP) – 2.2 (mNG) kDa) as the reporter of the sensor, which allows the system to conserve energy and resources significantly that leading more units of the reporter to be synthesized compared to synthesizing full-length sfGFP (230 amino acids long, 25.8 kDa) and mNG (227 amino acids long, 25.7 kDa) as the reporter. The overexpressed and purified large segment of the fluorescent protein (first-tenth β-strands (1-10)) would be supplemented into the cell-free reaction prior to activation of the sensor. The full-length sfGFP and mNG have previously been characterized in the CFS as bright reporter protein candidates.^9^ For the split mNG used in this study, we particularly utilized the complementation-enhanced version of the split mNG.^7^

In this study, we elucidated the feasibility of the split protein interaction compared to the whole-cell system and explored the complementation efficiencies and brightness between sfGFP and mNG split protein systems in the CFS. In addition, the gene ratios of the 1-10 and eleventh segments were investigated to enhance the output signal. Moreover, we improved the complementation of mNG split segments by adding SynZip peptide linkers^10^ and demonstrated the expansion of the cell-free biosensor toolbox with lower limits of detection.

## Results and Discussion

### Difference in complementation efficiencies across fluorescent protein and expression systems

The eleventh segments for both sfGFP and mNG were cloned into the pJL1 vector for cell-free reaction. For preliminary expression and fluorescence screening purposes, the 1-10 segments were also cloned into the same vector and co-expressed with the eleventh segment in the CFS to avoid potential low protein synthesis using pET vectors.^11^ For the whole-cell expression, the 1-10 segment genes were cloned in the pETBlue1 vector, ensuring their overexpression. Both full-length (unsplit) sfGFP and mNG were synthesized completely, generating the fluorescent in both cell-free and whole-cell expression systems (Fig. 1). When 1-10 segment were expressed in the CFS, sfGFP 1-10 segment exhibited complete fluorescence compared to its cell-free synthesized full-length counterpart, while the mNG 1-10 segment did not develop the fluorescence (Fig. 1). This phenomenon complicated the analysis of split sfGFP complementation in the CFS, especially since introducing the eleventh segment resulted in decreased fluorescence than both the 1-10 segments and full-length proteins (Fig. 1b, black bar). In contrast, the mNG 1-10 segment alone did not develop fluorescence on its own in the CFS. However, mNG did develop a small fluorescence increase in the presence of the eleventh segment (co-expression of 1-10 and eleventh segments) (Fig. 1a, black bar). To investigate whether the inherent fluorescence of the sfGFP 1-10 segment resulted from its expression in the CFS, we transitioned the split expression to a whole cell expression system. In the whole-cell system, neither the 1-10 segment alone nor its co-expression with the eleventh segment exhibited fluorescence, indicating the split mNG system is better aligned with the CFS (Fig. 1). The difference in fluorescence between the 1-10 segments of sfGFP and mNG could be attributed to sfGFP being a weak dimer, whereas mNG exists as a monomer.

**Figure 1.**
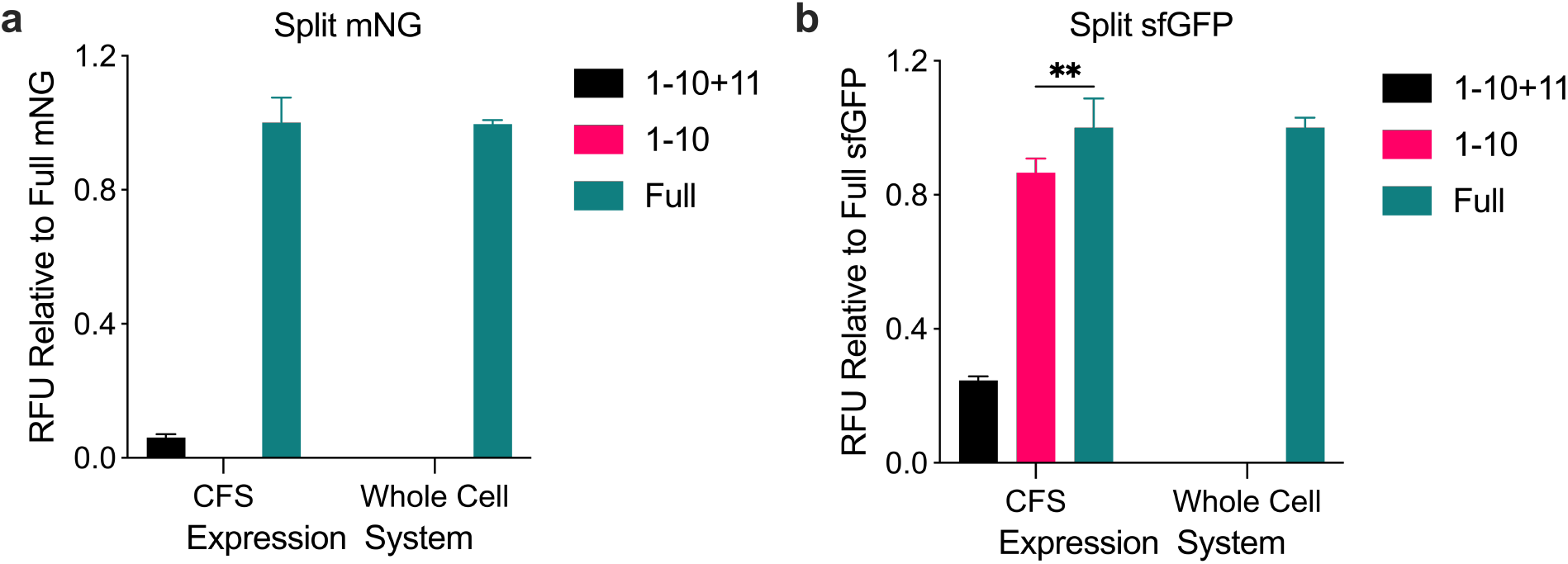
Split fluorescent protein expression in two protein expression systems. a) Split mNG expression in the CFS and whole-cell culture compared to full-length mNG. b) Split sfGFP expression in the CFS and whole-cell culture compared to full-length sfGFP. Black bar: co-expression of the 1-10 and the eleventh segments, Red bar: expression of the 1-10 segment alone, Green bar: full-length protein expression. Values represented as mean ± SD, n=3, **p<0.01.

### Varying gene concentrations of the split protein segments

Next, we investigated the gene concentrations of the split protein segments. The DNA concentrations in the CFS were varied to determine two objectives: 1) to ascertain the optimal ratio between the 1-10 segment gene and the eleventh segment gene for the effective functioning of the split mNG system, and 2) to differentiate whether the eleventh segment of the sfGFP was interfering with the folding of 1-10 segment of sfGFP or facilitating its assembly. We found that an equimolar ratio between 1-10 and the eleventh segments gene was optimal for the split mNG system. However, the equimolar ratio did not enhance the brightness up to the level of the full-length protein (data normalized to full-length mNG) (Fig. 2a). Interestingly, for split sfGFP system, a decrement in the eleventh segment gene concentration led to an upward in fluorescent output. This trend could imply that the eleventh segment might be impeding the expression of the 1-10 segment rather than aiding in its assembly (Fig. 2b).

**Figure 2.**
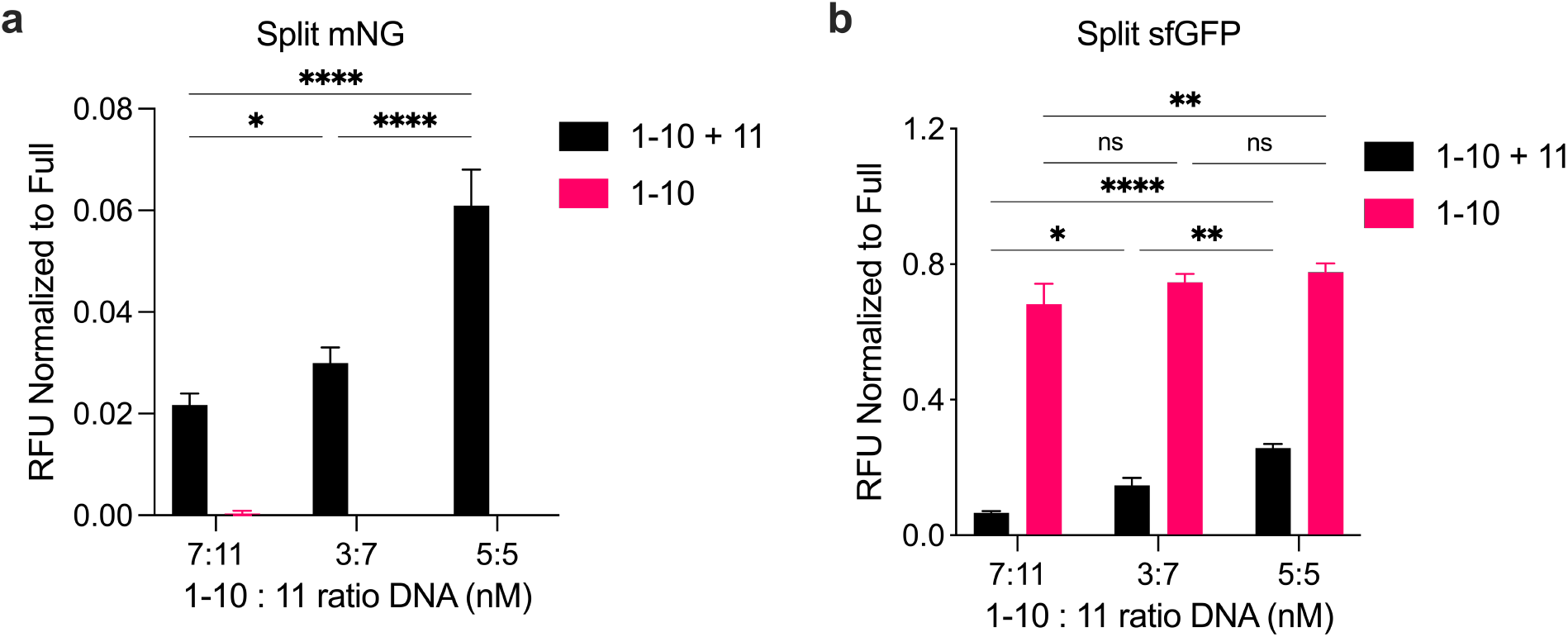
Varying DNA concentration ratio of the two split segments in the CFS. a) Split mNG expression values normalized to full-length mNG. b) Split sfGFP expression values normalized to full-length sfGFP. Black bar: 1-10 segment DNA ratio, Red bar: eleventh segment DNA ratio. Values represented as mean ± SD, n=3, *p<0.05, **p<0.01, ****p<0.0001.

### Improving split mNG interaction with SynZip peptide linkers

To enhance the conjoining of the eleventh and large 1-10 segments, we incorporated linker peptides known as SynZip 17 and 18 on each respective segment. These SynZip linkers have previously demonstrated their efficacy in improving split T7 RNA polymerase assemblies,^12^ and we observed similar enhancement in this study. The SynZip 17 sequence (Table S1) was added to the C-terminus of the mNG 1-10 segment gene in the pETBlue1 vector. The large mNG 1-10 segment was then expressed in *Escherichia coli* strain BL21(DE3) star to facilitate protein purification using the N-terminal histidine tag (Figure S1). The purified large mNG segment was supplied to the CFS. In contrast, the SynZip 18 sequence (Table S1) was integrated at the N-terminus of the eleventh segment in the pJL1 vector and expressed in the CFS, increasing the peptide size from 2.2 kDa to 7.3 kDa. The amount of purified 1-10 segment was varied, along with the DNA concentration of the eleventh segment in the CFS (Fig. 3a). As the DNA concentration of the eleventh segment decreased and purified 1-10 segment concentration increased, we achieved a fluorescence signal representing 73.6% ± 0.2% of the full-length mNG under the same conditions. However, it was noticed that the addition of the purified 1-10 segment did inhibit protein expression of the reaction overall, leading to a decrease of 28.0% ± 1% at 3 nM DNA concentration of full-length mNG and 66.3% ± 1.5% decrease at 1 nM (Fig. 3b). Potential causes for this reduction in expression could be the trace proteins in the purified product or the possibility that the isolated 1-10 segment may aggregate in the absence of its eleventh segment counterpart, thereby interfering with the stoichiometric balance of the other proteins in the CFS.^3,5^ For example, as illustrated in Figure 3b, a decrease in DNA concentration results in less excess mRNA in the system, which might amplify purified 1-10 segment’s toxicity to the transcriptional machinery. Nevertheless, the addition of SynZip linkers enhanced the fluorescence of the conjoined split mNG, elevating it by 4.8 ± 0.2-fold in the CFS and allowing for fluorescence to occur in general in the whole-cell expression system, surpassing the performance in the CFS (Fig. 3c).

**Figure 3.**
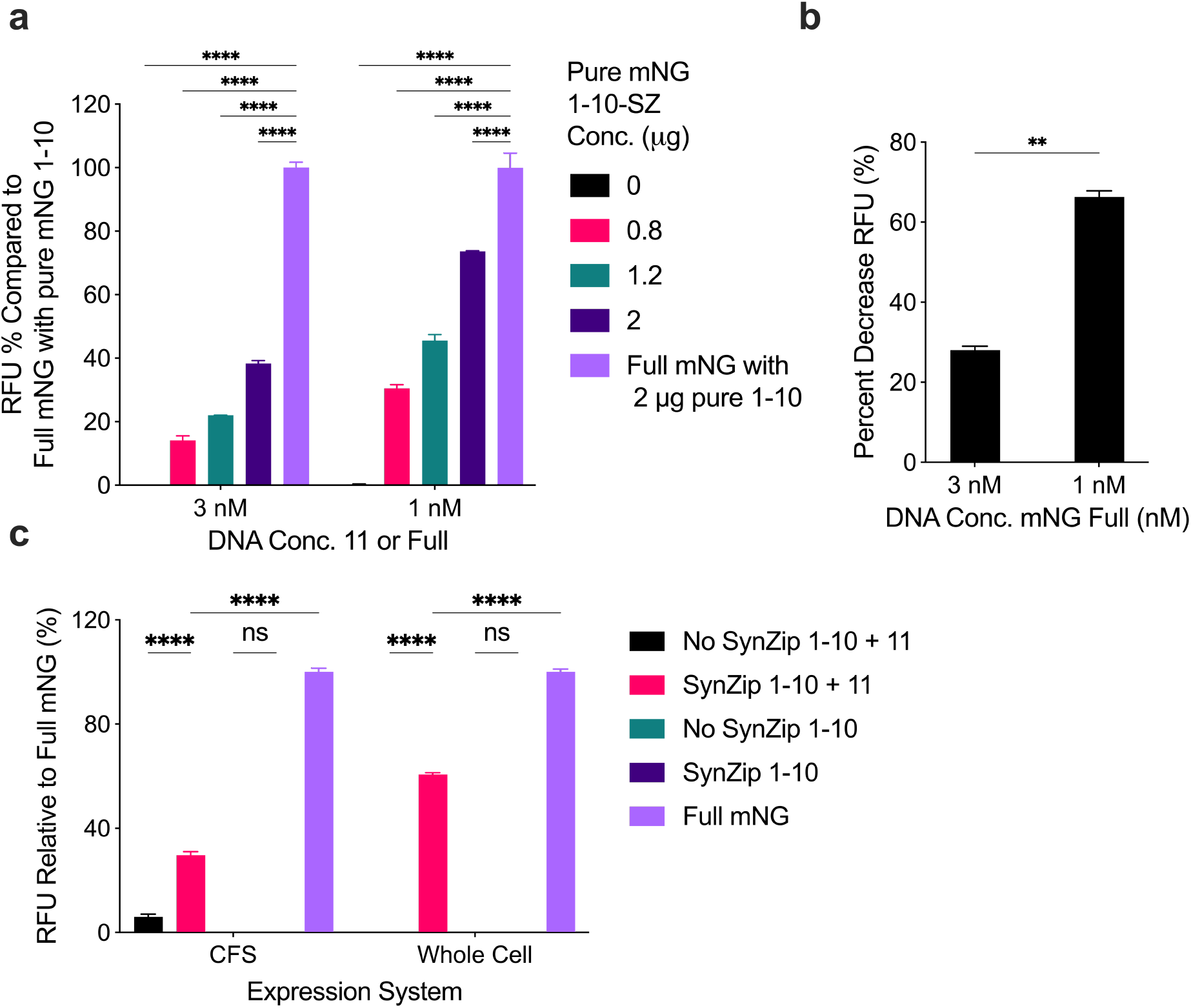
Improvement of split mNG with the addition of SynZip linkers. a) Percent fluorescent output compared to full-length mNG with purified mNG 1-10 segment in the reactions with varying concentrations of purified mNG 1-10 segment and mNG eleventh segment DNA. b) Percent decrease in protein expression when purified mNG 1-10 segment to the CFS expressing full-length mNG at varying concentrations. c) Split mNG expression in the CFS and whole-cell culture compared to full-length mNG with or without SynZip linkers. Values represented as mean ± SD, n=3, **p<0.01, ****p<0.0001.

## Conclusion

Here, we debuted the split fluorescent protein system in the CFS for the first time. The sfGFP 1-10 segment alone exhibited fluorescence in the CFS but remained inactive in a whole-cell system. This could potentially suggest that the cell-free environment supports the improved protein folding dynamics of the incomplete structure of the sfGFP 1-10 segment. However, this feature also proved that the split sfGFP system would not be a good candidate for cell-free biosensor design and development. In contrast, split mNG emerged as a promising toolbox for the CFS biosensor, given its contingent conjoining in the CFS and its non-fluorescence in the absence of the eleventh segment. Yet, it is noteworthy that even upon optimization of the gene concentration ratio in the CFS, its fluorescence was substantially diminished to only 6% of the full-length mNG. Aiming to enhance this complementation, we employed SynZip peptide linkers – added to the termini of the two protein segments. The addition of the SynZip linkers resulted in a 73.6% fluorescence signal compared to the full-length mNG and a 4.8-fold increase in expression compared to the mNG split system without the SynZip linkers. However, a reduction in overall system activity was observed upon introducing the purified mNG 1-10 segment, resulting in a 28-66% decline in protein expression. In conclusion, the mNG split system with the SynZip peptide linkers developed in this study has the substantial potential to serve as a robust reporter for the cell-free biosensors that are required to utilize more energy towards sensing low levels of analyte and less towards making the reporter protein itself.

## Methods

### Strains and plasmids

*Escherichia coli* strains Subcloning Efficiency™ *DH5α* [Genotype F^-^ Φ80*lac*ZΔM15Δ(*lac*ZYA-*arg*F) U169 *rec*A1 *end*A1 *hsd*R17(r_K_^-^, m_K_^+^) *pho*A *sup*E44 *thi*-1 *gyr*A96 *rel*A1 λ-] and BL21(DE3) star [Genotype F^-^ *ompT hsdS*_B_ (r_B_^-^ m_B_^-^) *gal dcm rne131* (DE3)] were used for cloning and a source of the cell extract, respectively (Invitrogen, Waltham, MA). The *E. coli* cells were grown in either Luria–Bertani (LB) media (10 g/L of tryptone, 5 g/L of yeast extract, 10 g/L of sodium chloride in Milli-Q water) or 2xYTPG media (16 g/L of tryptone, 10 g/L of yeast extract, 5 g/L of sodium chloride, 7 g/L of potassium phosphate dibasic, 3 g/L of potassium phosphate monobasic, pH 7.2, and 0.1 M of glucose in Milli-Q water).

Plasmids were assembled using the Gibson assembly. Split mNG genes were obtained from the Addgene. The 1-10 segment sequence was obtained from pSFFV_mNG3K(1-10) (Plasmid #157993)^7^ and inserted into pETBlue1 with a N-terminus histidine tag. The eleventh segment was taken from pET_mNG2(1-10)_32aalinker_mNG2(11) (Plasmid #82611)^6^ and inserted into pJL1. Split sfGFP was created by splitting the eleventh β-strand sequence in pJL1-sfGFP from the 1-10 segments and inserting it into pJL1 vector. The 1-10 segment sequence was inserted into pETBlue1 with a N-terminus histidine tag. SynZip linker sequences were later added by inserting the respective genes into pETBlue1 and pJL1 vectors with SynZip linkers already present. Plasmids were transformed into DH5α electrocompetent cells for cloning and plasmid purification (Qiagen Plasmid Midi Kit) for sequencing and cell-free reaction.

### Whole-cell protein expression and fluorescent measurement

The pETBlue1 plasmids containing the 1-10 and the eleventh segment genes and full-length sfGFP and mNG in pJL1 were transformed into *E. coli* BL21(DE3) star competent cells via electroporation. A colony (BL21, 1-10 segment harboring plasmid) was selected and grown overnight in a 5 mL LB for the second transformation of the plasmid harboring the eleventh segment, while the other colonies were saved for 1-10 segment-only overexpression. The next day, the cells were harvested and washed in a series of 80% glycerol solution (4 °C, 5,000 rpm, 5 min). The OD_600_ of the final competent cell was determined to be 0.8-1.0. 50 μL of the competent cell was used to transform the eleventh segment harboring plasmid pJL1. The cells were selected under carbenicillin and kanamycin antibiotics, working concentration at both 50 μg/mL. Three green fluorescent protein gene and segment harboring cells (full-length, 1-10 segment only, 1-10 with the eleventh segments) were cultured overnight in 5 mL LB. The overnight cultures were then inoculated to 50 mL LB in a 1:500 ratio. At OD_600_ 0.5, the cells were induced with 1mM IPTG. The whole-cell fluorescence was read when the OD600 reached 3.0 with a Synergy HTX multi-mode microplate reader (BioTek, Winooski, VT, USA). Excitation and emission wavelengths were 485 nm and 528 nm, respectively.

### Cell extract preparation and cell-free protein synthesis

Cell extract was prepared as described previously.^13,14^ Briefly, overnight cultured *E. coli* BL21(DE3) star in LB media was inoculated to sterilized 1 L 2xYTPG media in a 2.5-L baffled Tunair shake flask and the cells were cultured at 37 °C with vigorous shaking at 250 rpm. T7 RNA Polymerase expression was induced at OD 0.5 with 1 mM of isopropyl β-D-1-thiogalactopyranoside (IPTG). Cells were harvested at the mid-exponential phase of OD_600_ at 3.0. Cells were harvested (4 °C, 5,000 rpm, 10 min), and washed and resuspended in Buffers A and B, and lysed using sonication following the previous study.^9^ The lysate was centrifuged at 12,000 x g in 4 °C for 10 min and the supernatant was collected as cell extract. The cell extract was aliquoted and flash-frozen in liquid nitrogen and stored at a -80 °C freezer until use.

CFS components, including *E. coli* total tRNA mixture (from strain MRE600) ATP, GTP, CTP, UTP, Phosphoenolpyruvate, 20 amino acids, and other materials were purchased from Sigma-Aldrich (St. Louis, MO), Alfa Aesar (Haverhill, MA), and Fisher Scientific (Hampton, NH). Cell-free protein synthesis reactions were carried out, as mentioned previously.^9^ The reaction volume was 15 μL with the following components: 1.2 mM ATP; 0.85 mM each of GTP, UTP and CTP; 34.0 μg/mL L-5-formyl-5, 6, 7, 8-tetrahydrofolic acid (folinic acid); 170.0 μg/mL of *E. coli* tRNA mixture; 130 mM potassium glutamate; 10 mM ammonium glutamate; 12 mM magnesium glutamate; 2 mM each of 20 amino acids; 57 mM of HEPES buffer pH 7.2; 0.4 mM of nicotinamide adenine dinucleotide (NAD); 0.27 mM of coenzyme A; 4 mM of sodium oxalate; 1 mM of putrescine; 1.5 mM of spermidine; 33 mM phosphoenolpyruvate (PEP); and 27% v/v of cell extract. Plasmid DNA was added at a concentration of 8 nM unless otherwise noted. Cell-free protein synthesis reactions were carried out for 20 hours to ensure completion at 30 °C.

### Protein Purification

A colony was selected and grown overnight at 37 °C in 5 mL LB media with shaking at 250 rpm. The next day, the culture was inoculated into a larger LB media culture of 500 mL at a 1:500 ratio. The cells were then cultured at 37 °C with shaking at 300 rpm in a 2.5 mL Tunair shake flask. Once the culture reached OD_600_ 0.5, 1 mM final concentration of IPTG was added to the culture to induce T7 RNA polymerase expression and subsequent expression of 1-10 segment. The culture flask was then moved to a 20 °C incubator (250 rpm), and the cells were cultured overnight and harvested in the morning. His-tag purification was performed using the harvested cell pellets and the Qiagen Ni-NTA resin, followed by the manufacturer’s protocol. The resin was washed five times with 40 mM imidazole in the wash buffer and eluted with 250 mM imidazole in the elution buffer. The eluate was concentrated with 10 kDa MWCO Amicon centrifugal filters. The buffer was replaced with a storage buffer compatible with the cell-free reaction. Storage buffer consisted of 20 mM sodium phosphate, pH 7.7, 1 mM EDTA, 1 mM DTT, 5% glycerol, and 100 mM NaCl. The purified protein was then aliquoted to small aliquots of 25 uL and flash-frozen in liquid nitrogen. This storage method retained the activity of the purified protein. Once the sample was thawed, it was not re-frozen and saved to use again to retain the activity.

### Protein analysis

The relative fluorescence units (RFU) of the synthesized fluorescent proteins were measured by the multi-well plate fluorometer (Synergy HTX, BioTek, Winooski, VT). 5 μL of the cell-free synthesized fluorescent protein and 45 μL of Milli-Q water were mixed in a 96-well half-area black plate (Corning Incorporated, Corning, NY). The plate was mixed in the plated reader orbitally at medium speed for 15 s and read at the height of 1.5 mm with a gain of 50. The excitation and emission spectra are 485 and 528 nm, respectively. The cell-free synthesized protein was visualized by Coomassie blue staining after protein gel electrophoresis using pre-casted 4-12% Bis-Tris gradient gel (Invitrogen, Waltham, MA). Purified protein concentration was measured by Bradford Assay.

### Statistical analysis

Statistical analyses were conducted using GraphPad Prism 8.4.3 (GraphPad Software) with a 5% significance level. For the parametric analysis of data from the quantification of the synthesized protein, two-way ANOVA followed by Dunnett’s test was used.

## Conflict of interest

The authors declare that the research was conducted in the absence of any commercial or financial relationships that could be construed as a potential conflict of interest.

## Author contribution

C.E.C. and Y.-C.K. conceived the project. C.E.C., C.J.H., and Y.-C.K. designed and conceptualized experiments. C.E.C., C.J.H., K.D.D., and S.C.L. prepared and performed experiments and acquired data. C.E.C. and C.J.H. analyzed and interpreted data. C.E.C. wrote the original manuscript. C.E.C., C.J.H., and Y.-C.K. revised and edited the manuscript. All authors contributed to the article and approved the submitted version. C.E.C. and C.J.H. contributed to this work equally.

## Acknowledgment

This work was supported by the National Science Foundation (Award No. 2223720) and the USDA National Institute of Food and Agriculture (HATCH, Accession No. 1021535, Project No. LAB94414).

## Notes

### Competing Interest Statement

The authors have declared no competing interest.

