## Supporting Information for "Expanding the Cell-Free Reporter Protein Toolbox by Employing a Split mNeonGreen System to Reduce Protein Synthesis Workload"

[†] These authors contributed equally.

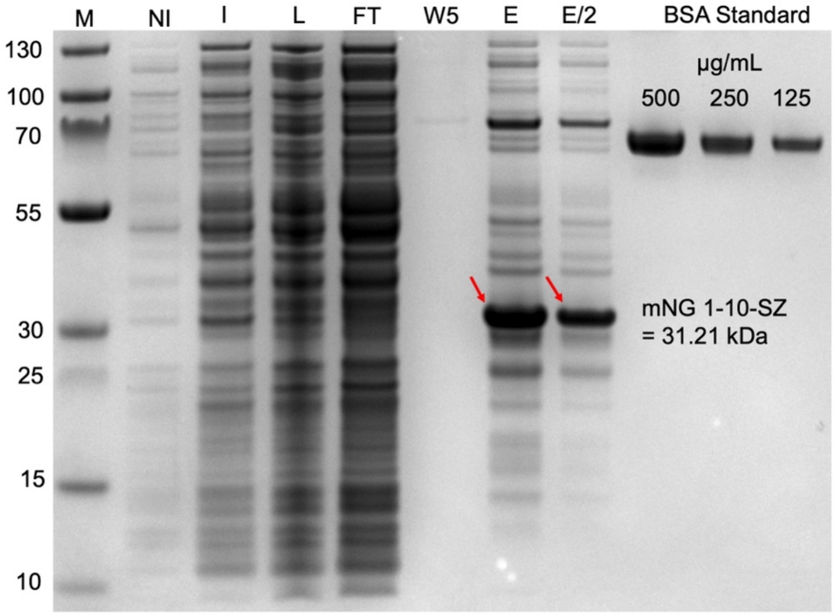


**Figure S1.** Purified N-His_mNG_1-10_SynZip (31.21 kDa). Lane 1: Protein marker, lane 2: non-induced cell culture sample, lane 3: induced cell culture sample, lane 4: lysate of harvested sample, lane 5: flow-through, lane 6: last wash (5th), lane 7: elution, lane 8: elution diluted by half volume, lane 9: BSA standard 500 µg/mL, lane 10: BSA standard 250 µg/mL, lane 11: BSA standard 125 µg/mL.

**Table S1.** DNA sequences (5' - 3') used in this study.

| | |
|---|---|
| *SynZip17 + linker* | agcatcgcggcgaccctggagaacgatctggcgcgtctggaaaacgaaaacgctcgtttggaaaaagacatcgcgaacctggaacgtgacctggcgaaactggagcgtgaagaagcgtacttcggcggtagtggtggca |
| *SynZip18 + linker* | agaacggaggttcaggtggtggatccaacgaaaaagaagaactgaaatccaaaaaagcggaactgcgcaaccgtatcgaacagctgaaacagaaacgtgaacaactgaagcagaaaatcgcgaacctgcgtaaagaaatcgaagcttacaaa |
| *sfGFP* | atgagcaaaggtgaagaactgtttaccggcgttgtgccgattctggtggaactggatggcgatgtgaacggtcacaaattcagcgtgcgtggtgaaggtgaaggcgatgccacgattggcaaactgacgctgaaatttatctgcaccaccggcaaactgccggtgccgtggccgacgctggtgaccaccctgacctatggcgttcagtgttttagtcgctatccggatcacatgaaacgtcacgatttctttaaatctgcaatgccggaaggctatgtgcaggaacgtacgattagctttaaagatgatggcaaatataaaacgcgcgccgttgtgaaatttgaaggcgataccctggtgaaccgcattgaactgaaaggcacggattttaaagaagatggcaatatcctgggccataaactggaatacaactttaatagccataatgtttatattacggcggataaacagaaaaatggcatcaaagcgaattttaccgttcgccataacgttgaagatggcagtgtgcagctggcagatcattatcagcagaataccccgattggtgatggtccggtgctgctgccggataatcattatctgagcacgcagaccgttctgtctaaagatccgaacgaaaaacgggaccacatggttctgcacgaatatgtgaatgcggcaggtattacgtggagccatccgcagttcgaaaaataa |
| *sfGFP 1-10 segment* | atgcatcatcaccatcaccacattgaagatggccgtagcaaaggtgaagaactgtttaccggcgttgtgccgattctggtggaactggatggcgatgtgaacggtcacaaattcagcgtgcgtggtgaaggtgaaggcgatgccacgattggcaaactgacgctgaaatttatctgcaccaccggcaaactgccggtgccgtggccgacgctggtgaccaccctgacctatggcgttcagtgttttagtcgctatccggatcacatgaaacgtcacgatttctttaaatctgcaatgccggaaggctatgtgcaggaacgtacgattagctttaaagatgatggcaaatataaaacgcgcgccgttgtgaaatttgaaggcgataccctggtgaaccgcattgaactgaaaggcacggattttaaagaagatggcaatatcctgggccataaactggaatacaactttaatagccataatgtttatattacggcggataaacagaaaaatggcatcaaagcgaattttaccgttcgccataacgttgaagatggcagtgtgcagctggcagatcattatcagcagaataccccgattggtgatggtccggtgctgctgccggataatcattatctgagcacgcagaccgttctgtctaaagatccgaacgaaaaataa |
| *sfGFP 11th segment* | atgcgggaccacatggttctgcacgaatatgtgaatgcggcaggtattacgtaa |
| *mNeonGreen (E. coli codon optimized)* | atggcaagtctacccgctacacacgaattacacatcttcggtagtattaacggggtggattttgatatggttggtcagggtactggaaacccgaatgacggctatgaggaactgaacctgaagtccaccaaaggtgatctgcaattctctccgtggattctggttccgcatatcggttacggcttccatcaatatttaccgtatccagatggcatgagcccatttcaggcggccatggtcgacggctctggttaccaagtgcatagaaccatgcagttcgaggacggcgcgagcctgacggtgaactaccgctacacctacgagggttcccacatcaaaggcgaagcgcaggtgaaaggtactggcttcccggcagacggtccggttatgaccaatagcttgaccgcggctgactggtgccgttcgaagaagacgtacccgaatgacaaaaccattatctccaccttcaagtggagctataccaccggcaacggtaaacgttaccgcagcactgcgcgtaccacatacaccttcgccaaaccgatggcagctaattatttgaagaaccagccgatgtatgtctttcgtaaaacggagcttaagcacagcaagaccgagctcaactttaaagaatggcaaaaggcgtttaccgatgttatgggtatggatgaactgtataaatggtcccatccgcagtttgaaaagtaa |
| *mNeonGreen 1-10 segment* | atgcatcatcaccatcaccacattgaagatggccgtgtgagcaagggtgaggaggataacatggcctctctcccagcgactcatgagttacacatctttggctccatcaacgatgtggactttgacatggtgggtcagggtaccggcaatccaaatgaaggttatgaggagttaaacctgaagtccaccaagggcgacctccagttctccccctggattctggtccctcatatcgggtatggcttccatcagtacctgccctaccctgacgggatgtcgcctttccaggccgccatggtagatggctccggataccaagtccatcgcacaatgcagtttgaagatggcgcctcccttactgttaactaccgctacacctacgagggaagccacatcaaaggagaggcccaggtgatagggactggtttccctgctgacggtcctgtgatgaccaacacgctgaccgctgcggactggtgcatgtcgaagatgacttaccccaacgacaaaaccatcatcagtacctttaagtggagttacatcactgtaaat |

| | |
|---|---|
| | ggcaaacgctaccggagcactgcgcggaccacctacacctttgccaagccaatggcggctaactatctgaagaaccagccgatgtacgtg<br>ttccgtaagacggagctcaagcactccatgtaa |
| *mNeonGreen 11$^{th}$ segment* | atgaccgagctcaacttcaaggagtggcaaaaggcctttaccgatatgatgtaa |
| *mNeonGreen 1-10 segment + SynZip 17* | atgcatcatcaccatcaccacattgaagatggccgtgtgagcaagggtgaggaggataacatggcctctctcccagcgactcatgagttac<br>acatctttggctccatcaacgatgtggactttgacatggtgggtcagggtaccggcaatccaaatgaaggttatgaggagttaaacctgaagt<br>ccaccaagggcgacctccagttctccccctggattctggtccctcatatcgggtatggcttccatcagtacctgccctaccctgacgggatgt<br>cgcctttccaggccgccatggtagatggctccggataccaagtccatcgcacaatgcagtttgaagatggcgcctcccttactgttaactacc<br>gctacacctacgagggaagccacatcaaaggagaggcccaggtgatagggactggtttccctgctgacggtcctgtgatgaccaacacg<br>ctgaccgctgcggactggtgcatgtcgaagatgacttaccccaacgacaaaaccatcatcagtacctttaagtggagttacatcactgtaaat<br>ggcaaacgctaccggagcactgcgcggaccacctacacctttgccaagccaatggcggctaactatctgaagaaccagccgatgtacgtg<br>ttccgtaagacggagctcaagcactccatggagaacggaggttcaggtggtggatccaacgaaaaagaagaactgaaatccaaaaaagc<br>ggaactgcgcaaccgtatcgaacagctgaaacagaaacgtgaacaactgaagcagaaaatcgcgaacctgcgtaaagaaatcgaagctt<br>acaaataa |
| *mNeonGreen 11$^{th}$ segment + SynZip 18* | atgagcatcgcggcgaccctggagaacgatctggcgcgtctggaaaacgaaaacgctcgtttggaaaaagacatcgcgaacctggaacg<br>tgacctggcgaaactggagcgtgaagaagcgtacttcggcggtagtggtggcaagaccgagctcaacttcaaggagtggcaaaaggcct<br>ttaccgatatgatgtaa |
